# Broad-spectrum antiviral activity of clinically approved CYP3A inhibitors against pathogenic human coronaviruses in vitro

**DOI:** 10.1101/2023.08.23.554463

**Authors:** Lara Gallucci, James Bazire, Andrew D Davidson, Iart Luca Shytaj

## Abstract

Coronaviruses pose a permanent risk of outbreaks, with three highly pathogenic species and strains (SARS-CoV, MERS-CoV, SARS-CoV-2) having emerged in the last twenty years. Limited antiviral therapies are currently available and their efficacy in randomized clinical trials enrolling SARS-CoV-2 patients has not been consistent, highlighting the need for more potent treatments. We previously showed that cobicistat, a clinically approved inhibitor of Cytochrome P450-3A (CYP3A), has direct antiviral activity against early circulating SARS-CoV-2 strains *in vitro* and in Syrian hamsters. Cobicistat is a derivative of ritonavir, which is co-administered as pharmacoenhancer with the SARS-CoV-2 protease inhibitor nirmatrelvir, to inhibit its metabolization by CPY3A and preserve its antiviral efficacy. Here, we used automated imaging and analysis for a screening and parallel comparison of the anti-coronavirus effects of cobicistat and ritonavir. Our data show that both drugs display antiviral activity at low micromolar concentrations against multiple SARS-CoV-2 variants *in vitro*, including epidemiologically relevant Omicron subvariants. Despite their close structural similarity, we found that cobicistat is more potent than ritonavir, as shown by significantly lower EC50 values in monotherapy and higher levels of viral suppression when used in combination with nirmatrelvir. Finally, we show that the antiviral activity of both cobicistat and ritonavir is maintained against other human coronaviruses, including HCoV-229E and the highly pathogenic MERS-CoV. Overall, our results demonstrate that cobicistat has more potent anti-coronavirus activity than ritonavir and suggest that dose adjustments could pave the way to the use of both drugs as broad-spectrum antivirals against highly pathogenic human coronaviruses.

## 1. Introduction

The severe acute respiratory syndrome coronavirus 2 (SARS-CoV-2) pandemic has underlined the need for broadly effective antivirals that can be quickly deployed during outbreaks. In this regard, the *Coronaviridae* family, to which SARS-CoV-2 belongs, represents a permanent threat due to the endemic persistence of its human-infecting species and strains and to the frequent zoonotic events leading to cross-species transmission to humans (V’Kovski et al., 2021). Coronaviruses are enveloped viruses bearing a positive, single stranded, RNA genome encoding four structural proteins, the spike (S), the envelope (E), the membrane (M) and the nucleoprotein (N), and at least three key enzymes, the RNA-dependent RNA polymerase (RdRp), the main protease (M^pro^) and the papain-like protease (PLpro) (V’Kovski et al., 2021). Apart from SARS-CoV-2, recent coronavirus outbreaks of high morbidity and mortality were caused by SARS-CoV and by the Middle East respiratory syndrome–related coronavirus (MERS-CoV). In addition, coronaviruses include four species (HCoV-229E, HCoV-OC43, HCoV-NL63, HCoV- HKU1) that are endemic but cause less severe symptoms, typically common cold and upper respiratory tract infections (Sariol & Perlman, 2020; V’Kovski et al., 2021).

A massive drug discovery effort has led to the identification of antiviral treatments against SARS-CoV-2, mainly repurposed from other indications (Edwards et al., 2022; Taibe et al., 2022). Due to the emergence of multiple SARS-CoV-2 variants of concern (VOCs) and variants of interest (VOIs) with different infectivity and morbidity, the spectrum of activity of these antiviral treatments has also been investigated. Treatments selectively targeting the association of the S protein to the cellular ACE-2 receptor, such as monoclonal antibodies, have shown significant loss of efficacy with the emergence of Omicron VOCs and VOIs displaying escape mutations on S protein epitopes (Cox et al., 2023). On the other hand, drugs targeting key viral enzymes such as the M^pro^ inhibitor nirmatrelvir and the RdRp inhibitors remdesivir and molnupiravir, have generally shown robust cross-variant efficacy even retaining antiviral activity against other pathogenic human coronaviruses. These include MERS-CoV (de Wit et al., 2020; Sheahan et al., 2017; Sheahan, Sims, Leist, et al., 2020; Sheahan, Sims, Zhou, et al., 2020), HCoV-229E, HCoV-OC43 (Brown et al., 2019; Li et al., 2023; Li et al., 2022; Liu et al., 2023) and HCoV-NL63 (Li et al., 2022; Liu et al., 2023), although the latter was recently reported to be partially resistant to nirmatrelvir (Li et al., 2023).

Despite these advances, clinically approved options for treating SARS-CoV-2 infection are still very limited. The most promising nucleoside analogues, remdesivir and molnupiravir, were not consistently associated with clinical benefit in randomized trials (Butler et al., 2023; Consortium, 2022). At least in the case of remdesivir, the discrepancy between pre-clinical and clinical results might be due to the unfavorable pharmacokinetics and rapid drug excretion *in vivo* (Leegwater et al., 2022) which could however be improved by a recently developed, orally available, derivative (Cao et al., 2023). The M^pro^ inhibitor nirmatrelvir is likewise rapidly metabolized *in vivo*, but its clearance can be significantly reduced through the co-administration of the cytochrome P450 3A (CYP3A) inhibitor ritonavir (Lamb, 2022). The co-formulation of nirmatrelvir with ritonavir (named Paxlovid) (Lamb, 2022) is currently the most widely approved antiviral treatment for non-hospitalized coronavirus disease -19 (COVID-19) patients and was shown to significantly decrease the likelihood of disease progression in randomized clinical trials (Hammond et al., 2022). However, Paxlovid might be unable to decrease hospitalization rates in all age cohorts infected with Omicron subvariants (Arbel et al., 2022), and viral rebound after therapy has been reported in a sizable subset of individuals (Pandit et al., 2023), suggesting that more potent viral suppression might be necessary to clear the infection.

Prior to its use as a pharmacoenhancer in Paxlovid, ritonavir had been developed as an antiretroviral drug to inhibit the human immunodeficiency virus (HIV) protease (Lea & Faulds, 1996). Due to the emergence of more potent HIV protease inhibitors, ritonavir has been progressively repositioned as a booster to other antiretroviral drugs. As antiretroviral booster, ritonavir has been partially superseded by its derivative, cobicistat, which is devoid of anti-HIV activity, but retains the ability to inhibit CYP3A (Xu et al., 2010). In a previous study, we found that cobicistat, used at concentrations that are well tolerated but higher than those required for CYP3A inhibition, unexpectedly inhibited the fusion and replication of SARS-CoV-2 *in vitro* and in Syrian hamsters (Shytaj et al., 2022). However, in that work we only investigated two early-circulating (Worobey et al., 2020) SARS-CoV-2 isolates (Ger/BavPat1/2020 and Munich/BavPat2-ChVir984-ChVir1017/2020) and we did not test whether the parent drug of cobicistat, ritonavir, could exert similar antiviral effects.

In the present work we use automated imaging and analysis to perform a high-throughput parallel screening of the *in vitro* antiviral effects of cobicistat and ritonavir against eight past and currently circulating SARS-CoV-2 VOCs and VOIs, as well as against HCoV-229E and MERS-CoV. Our data show that, although both drugs have broad spectrum anti-coronavirus activity at low micromolar concentrations, cobicistat is consistently more potent, as shown by more pronounced maximal inhibition of viral replication, lower EC50 values and higher combination sensitivity scores upon co-treatment with nirmatrelvir. Considering the previous pre-clinical and clinical experience with CYP3A inhibitors, our study suggests that dose-adjustments to standard administration protocols could facilitate the leveraging and use of CYP3A inhibitors, and particularly cobicistat, as first-line monotherapy or combination treatment against human coronaviruses.

## 2. Materials and Methods

### 2.1. Cell lines and drug treatments

Vero E6 cells modified to constitutively express TMPRSS2 (VTN cells) (Matsuyama et al., 2020) were provided by the NIBSC Research Reagent Repository, UK. Human liver epithelial Huh-7 cells were a kind gift from Professor Mark Harris (University of Leeds). Cells were cultured in Dulbecco’s Modified Eagle’s medium, containing 4.5 g/l D-glucose, and GlutaMAX™ (DMEM, Gibco™, ThermoFisher) supplemented with 1 mM sodium pyruvate (Sigma-Aldrich), 10% fetal bovine serum (FBS, Gibco™, ThermoFisher) and 1% penicillin-streptomycin (Pen/Strep) at 37 °C in a humidified incubator in 5% CO2. The day before infection, cells were seeded at 7-10LJ×LJ10^3^ per well in µClear 96-well Microplates (Greiner Bio-one). Immediately before infection cells were treated with serial dilutions of cobicistat (Santa Cruz Biotechnology), ritonavir (Sigma Aldrich) and/or nirmatrelvir (MedChemExpress) diluted in Modified Eagle’s medium (2% FBS and 1% Pen/Strep). DMSO, in which all drug stocks were initially prepared, was used as vehicle control.

### 2.2. Viral stocks and infection

The following stocks were used: 1) SARS-CoV-2 variants; a) Wuhan (REMRQ0001 isolated in April, 2020 as previously described (Daly et al., 2020), b) Alpha (hCoV- 19/England/204690005/2020; GISAID ID: EPI_ISL_693401), c) Beta (hCoV-19/England/205280030/2020; GISAID ID: EPI_ISL_770441), d) Gamma (hCoV-19/England/520336_B1_P0/2021, GISAID ID: EPI_ISL_2080492), e) Delta (GISAID ID: EPI_ISL_15250227, isolated as previously described (Erdmann M, 2022), f) Omicron BA.1 (isolated as previously described (Dejnirattisai et al., 2022)), g) Omicron BA.4, and h) Omicron XBB.1, (the Alpha, Beta, Gamma and Omicron VOCs were kindly provided by Professor Wendy Barclay, Imperial College, London and Professor Maria Zambon, UK Health Security Agency); 2) MERS-CoV (isolate HCoV-EMC/2012, GenBank accession number NC_019843.3, kindly provided by Professor Fouchier, Erasmus Medical Center, Rotterdam, The Netherlands), and 3) HCoV229E (GenBank accession number NC_002645, kindly provided by Professor Stuart Siddell, University of Bristol). The genome sequences of all viruses were verified after stock production by Illumina sequencing. The infectious titer of each of the stock viruses was determined by serial dilution on VTN or Huh-7 cells (HCoV-229E) followed by detection of infected cells by immunofluorescence assay (at 6h – 8h post-infection) and automated image analysis as described below. Following drug treatment cells were infected at 0.05 MOI and kept for 24h at 37 °C in a humidified incubator in 5% CO2.

### 2.3. Immunofluorescence staining and image acquisition

One day post-infection, cells were fixed in 4 % paraformaldehyde (PFA) in PBS for 1h and then permeabilized with 0.1% Triton-X100 in PBS and blocked with 1% (w/v) bovine serum albumin. Cells were then stained using a monoclonal antibody targeting the SARS-CoV-2 N protein (1:1000 dilution; 200-401-A50, Rockland), the MERS-CoV N protein (1:500 dilution; 40068- RP01, Sino Biological) or double stranded-RNA (dsRNA) (1:500 dilution, J2 10010200, Scicons). After 45 minutes incubation, cells were stained with Alexa Fluor-conjugated secondary antibodies (1:3000 dilution, Invitrogen™, ThermoFisher) and DAPI (1:2000 dilution; D3571, Invitrogen). To determine the percentage of infected cells, images were acquired on an ImageXpress Pico Automated Cell Imaging System (Molecular Devices) using a 10X objective. Images encompassing the central 50% of the well were analyzed using a ImageXpress Pico Automated Cell Imaging System software (Molecular Devices). Representative images were prepared using Fiji, applying a macro to all images to ensure the same display adjustments across treatment conditions.

### 2.4. MTT assay

The effect of drug treatments on cell viability was assessed using the MTT [3-(4,5- dimethylthiazol-2-yl)-2,5-diphenyl tetrazolium bromide] assay (the MTT powder was obtained from Sigma Aldrich). VTN or Huh-7 cells were seeded the day before treatment at a concentration of 7L×L10^3^ cells per well in DMEM supplemented with 10% FBS and 1% Pen/Strep. The next day, cells were treated with vehicle (DMSO) or with cobicistat, ritonavir and/or nirmatrelvir diluted in MEM (supplemented with 2% FBS and 1% Pen/Strep) for 24h. The medium was then removed and replaced with 100 μL per well of fresh medium. To each well, 15 μL of 5mg/mL MTT in water were then added. After 2-4h the reaction was stopped by adding 100 μL per well of 10% (w/v) SDS in water. The absorbance values were acquired using a GloMAX® Explorer microplate reader (Promega) at 600 nm. After subtracting blank absorbance, the viability of drug-treated cells was normalized to vehicle controls.

### 2.5. Data analysis

The percentage of infected cells (*i.e.* cells positive to the N nucleoprotein or to dsRNA) was calculated using the viral infectivity protocol analysis of the ImageXpress Pico Automated Cell Imaging System software. The percentage of infected cells was then used to calculate the inhibition of viral replication for each treatment condition according to the following formula:

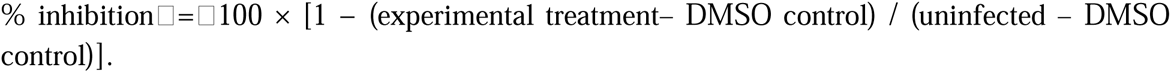

Half maximal effective concentrations (EC50) were calculated by non-linear regression and significant differences between the EC50s of cobicistat and ritonavir were assessed by extra sum of squares F-test. Drug interaction scores were evaluated using the web tool SynergyFinder plus (Zheng et al., 2022). Parameters analyzed included synergy (ZIP, Loewe, HSA, and Bliss models) and combination sensitivity (Malyutina et al., 2019). The relationship between maximal inhibition values and baseline infection was assessed by Spearman correlation, while pairwise comparisons were analyzed by paired *t*-test. All statistical analyses were conducted using GraphPad Prism v9.4.0 (GraphPad Software, San Diego, CA, USA).

## 3. Results

### 3.1. Antiviral activity of cobicistat and ritonavir against past and circulating SARS-CoV-2 VOCs and VOIs

To assess and compare the antiviral effects of the CYP3A inhibitors cobicistat and ritonavir we used a panel of eight SARS-CoV-2 variants including the reference Wuhan strain, four VOCs previously characterized by regional or global spread (Alpha, Beta, Delta, Gamma) and three recently or currently circulating Omicron VOCs and VOIs (*i.e.* Omicron BA.1, BA.4, XBB.1).

The experiments were conducted using Vero E6 cells constitutively expressing TMPRSS2 (*i.e.* VTN cells), which are highly permissive to SARS-CoV-2 infection and spread (Matsuyama et al., 2020).

Cells were pre-treated with a range (1.25-20 μM) of well-tolerated concentrations of cobicistat or ritonavir (Figure S1), infected for 24h, and stained by immunofluorescence for the specific (Figure S2A) expression of the viral N protein. The percentage of infected cells (*i.e.* cells positive for the N protein) was calculated through automated acquisition and analysis (as summarized in Figure S2B) and the effect of either drug was compared to the vehicle (DMSO) controls.

The results showed that, although both drugs displayed detectable antiviral activity against all tested variants, cobicistat was consistently more potent (Figures 1 and 2). The enhanced effects of cobicistat were evidenced by its lower EC50 values against all tested variants, which diverged from those of ritonavir with statistical significance in 5/8 variants (*i.e.* Wuhan, Alpha, Beta, Omicron BA.1 and Omicron BA.4) (Figure 1A-C and Figure 2B,C). Drug concentrations associated with antiviral activity were in the low-micromolar range, in line with our previous *in vitro* results testing cobicistat on the early SARS-CoV-2 isolate Ger/BavPat1/2020 (Shytaj et al., 2022).

**Figure 1.**
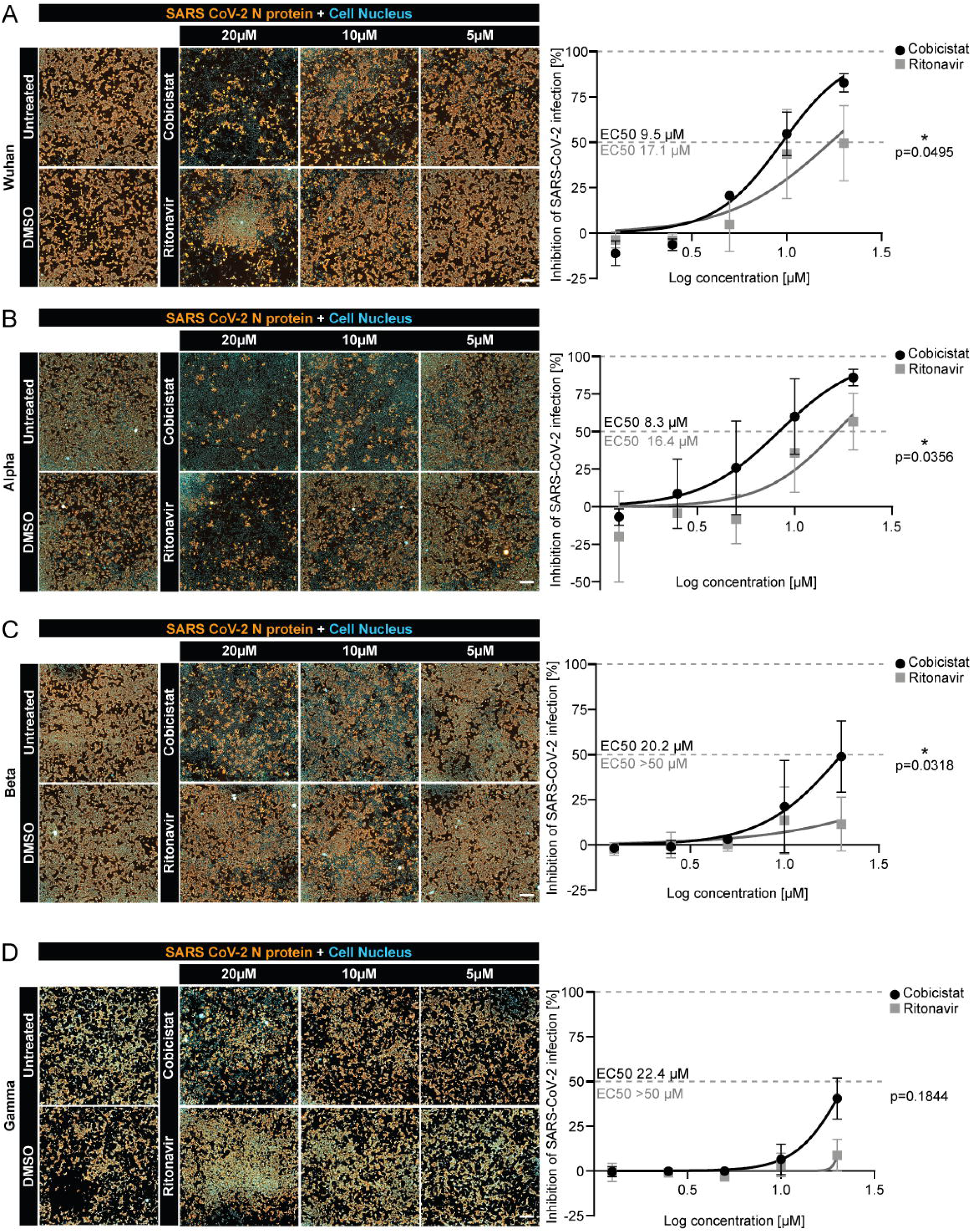
Efficacy of cobicistat and ritonavir against the reference Wuhan SARS-CoV-2 strain and regionally spread VOCs. Panels A-D. VTN cells were pre-treated with cobicistat or ritonavir at the indicated concentrations and then infected with the Wuhan (A), Alpha (B), Beta (C) or Gamma (D) variants. After 24h cells were fixed and stained for DAPI (blue) and the viral N protein (orange). Representative images are shown in the left panels. The proportion of cells positive for the N protein was calculated and used to derive the relative inhibition induced by each drug as compared to vehicle (DMSO) controls. Half maximal effective (EC50) concentrations of both cobicistat and ritonavir were calculated by non-linear regression using a variable Hill slope (right panels). The divergence of the curves used to fit EC50 values for cobicistat and ritonavir was assessed by extra sum-of-squares F test. Data points show mean ± SD of three independent experiments, each performed in duplicate. * p <0.05. Scale bar = 500 μm.

**Figure 2.**
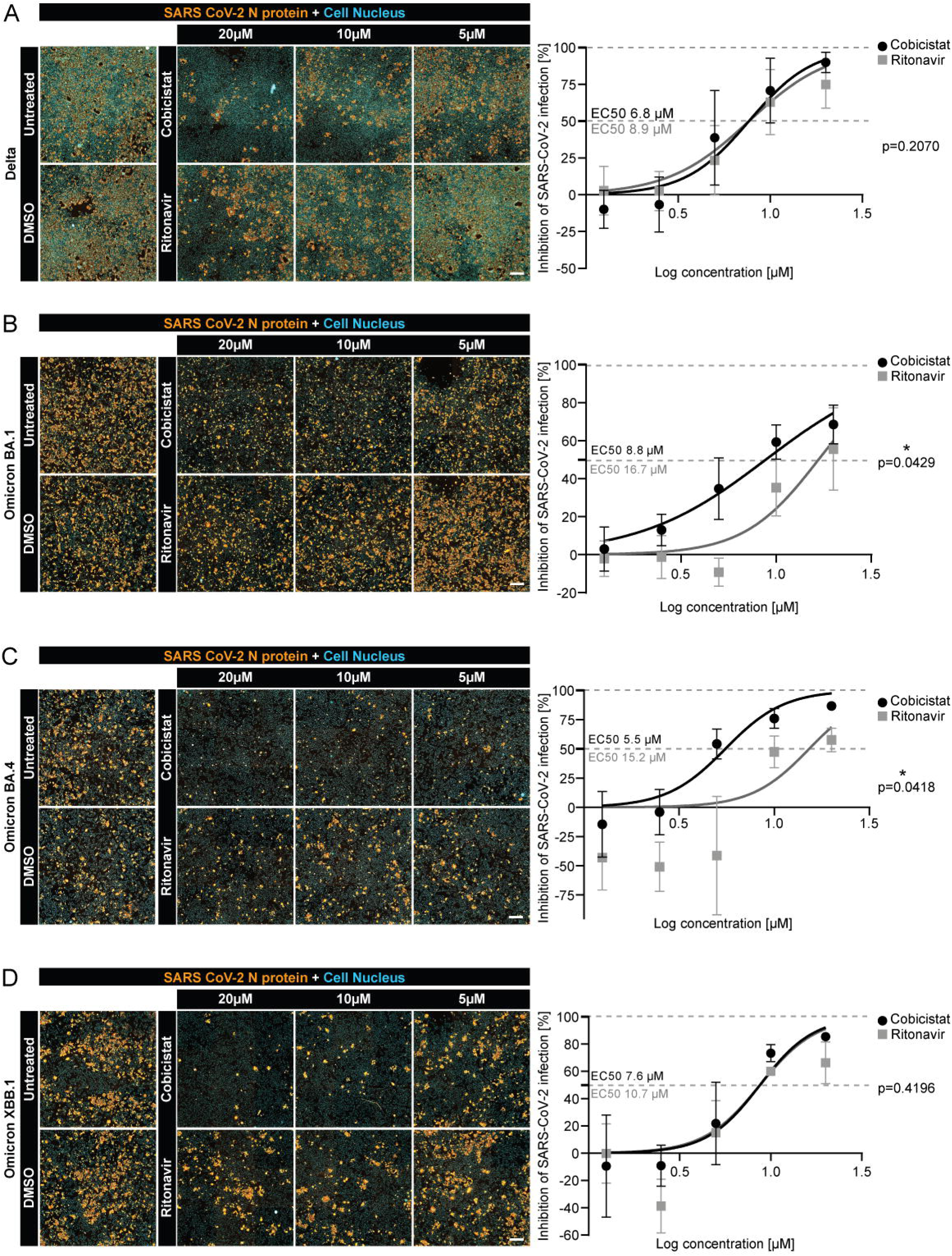
Efficacy of cobicistat and ritonavir against SARS-CoV-2 VOCs and VOIs characterized by current or previous global spread. Panels A-D. VTN cells were pre-treated with cobicistat or ritonavir at the indicated concentrations and then infected with the Delta (A), Omicron BA.1 (B), Omicron BA.4 (C) or Omicron XBB.1 (D) SARS-CoV-2 VOCs and VOIs. After 24h cells were fixed and stained for DAPI (blue) and the viral N protein (orange). Representative images are shown in the left panels. The proportion of cells positive for the N protein was calculated and used to derive the relative inhibition induced by each drug as compared to vehicle (DMSO) controls. Half maximal effective concentrations (EC50) of both cobicistat and ritonavir were calculated by non-linear regression using a variable Hill slope (right panels). The divergence of the curves used to fit EC50 values for cobicistat and ritonavir was assessed by extra sum-of-squares F test. Each data point shows mean ± SD of three independent experiments, each performed in duplicate. * p <0.05. Scale bar = 500 μm.

For each drug, differences in antiviral potency were also visible across variants, with high inhibitory effects obtained against the Alpha, Delta, Omicron BA.4 and Omicron XBB.1 variants (Figure 1B and Figure 2A,C,D), while the Beta and Gamma variants displayed the lowest susceptibility to inhibition (Figure 1C,D). Of note, the Beta and Gamma variants were also characterized by higher infectivity, with almost 100% of the cell culture being positive for the N protein in our DMSO controls. This suggested that the partial decrease in antiviral potency could be a consequence of the amount of baseline infection, rather than of variant-specific resistance. In line with this, when for each experiment maximum inhibition levels induced by either cobicistat or ritonavir were plotted with the corresponding baseline infection levels detected in the DMSO controls, an inverse correlation became evident (Figure 3). This correlation was highly significant for both cobicistat and ritonavir (p = 0.0014 and p <0.0001, respectively), although the extent of maximum inhibition induced by cobicistat was greater, corroborating its higher antiviral potency (Figure 3).

**Figure 3.**
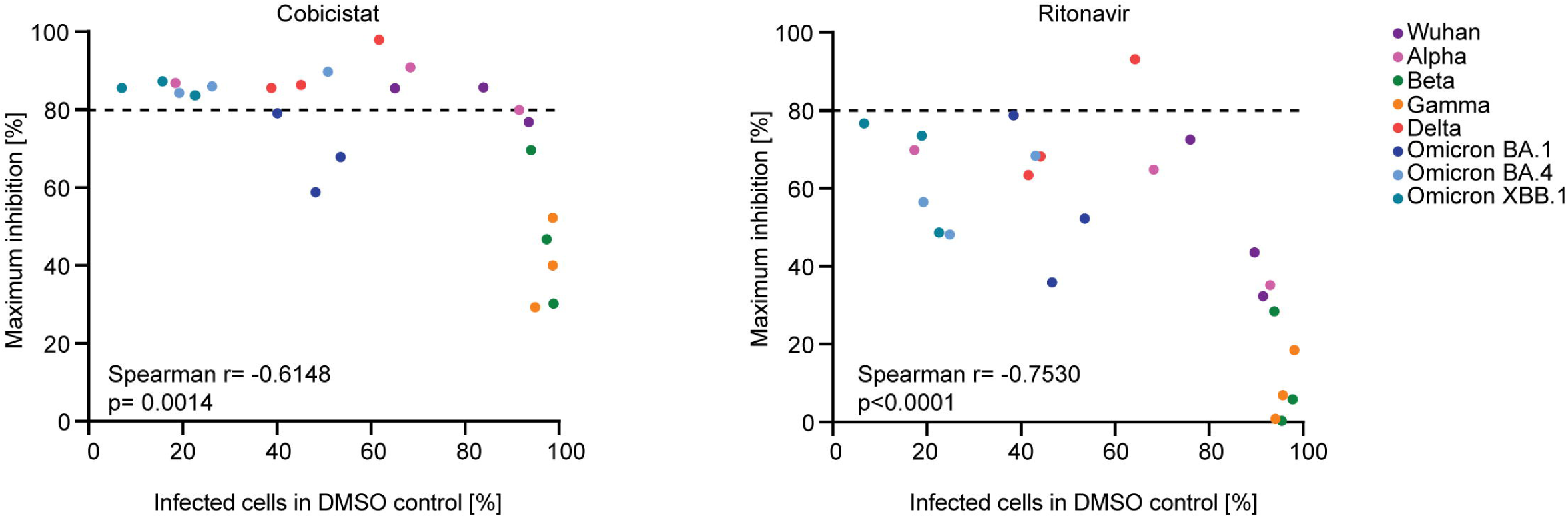
Correlation between antiviral efficacy of cobicistat or ritonavir and baseline SARS-CoV-2 infection. The data shown in Figures 1 and 2 were used to plot and correlate the maximum inhibitory effect of cobicistat or ritonavir against each SARS-CoV-2 variant (y axis) with the corresponding baseline percent of infected cells in vehicle (DMSO) treated controls (x axis). Each data point represents an experiment with a given SARS-CoV-2 variant, while the dotted line separates the experiments where a high level (*i.e.* >80%) of maximum inhibition was obtained. The statistical significance of the correlation was assessed by two-tailed Spearman nonparametric correlation.

Overall, these data show that both cobicistat and ritonavir can inhibit multiple SARS-CoV-2 VOCs and VOIs and that, although the strength of their antiviral activity is dependent on baseline infection levels, cobicistat is consistently more potent.

### 3.2. Higher SARS-CoV-2 inhibition through nirmatrelvir/cobicistat as compared to nirmatrelvir/ritonavir treatment

We then compared the antiviral effects obtained by combining cobicistat or ritonavir with the M^pro^ inhibitor nirmatrelvir, which is approved for clinical co-administration with ritonavir in the Paxlovid formulation (Lamb, 2022). The addition of nirmatrelvir to either cobicistat or ritonavir did not induce cytotoxic effects in VTN cells, even at the highest concentrations tested (Figure S3). In line with the effects previously described for Paxlovid, each drug combination significantly increased the antiviral effects of any individual compound for all tested SARS CoV-2 VOCs and VOIs (Figure 4A-D and Figure 5A-D).

**Figure 4.**
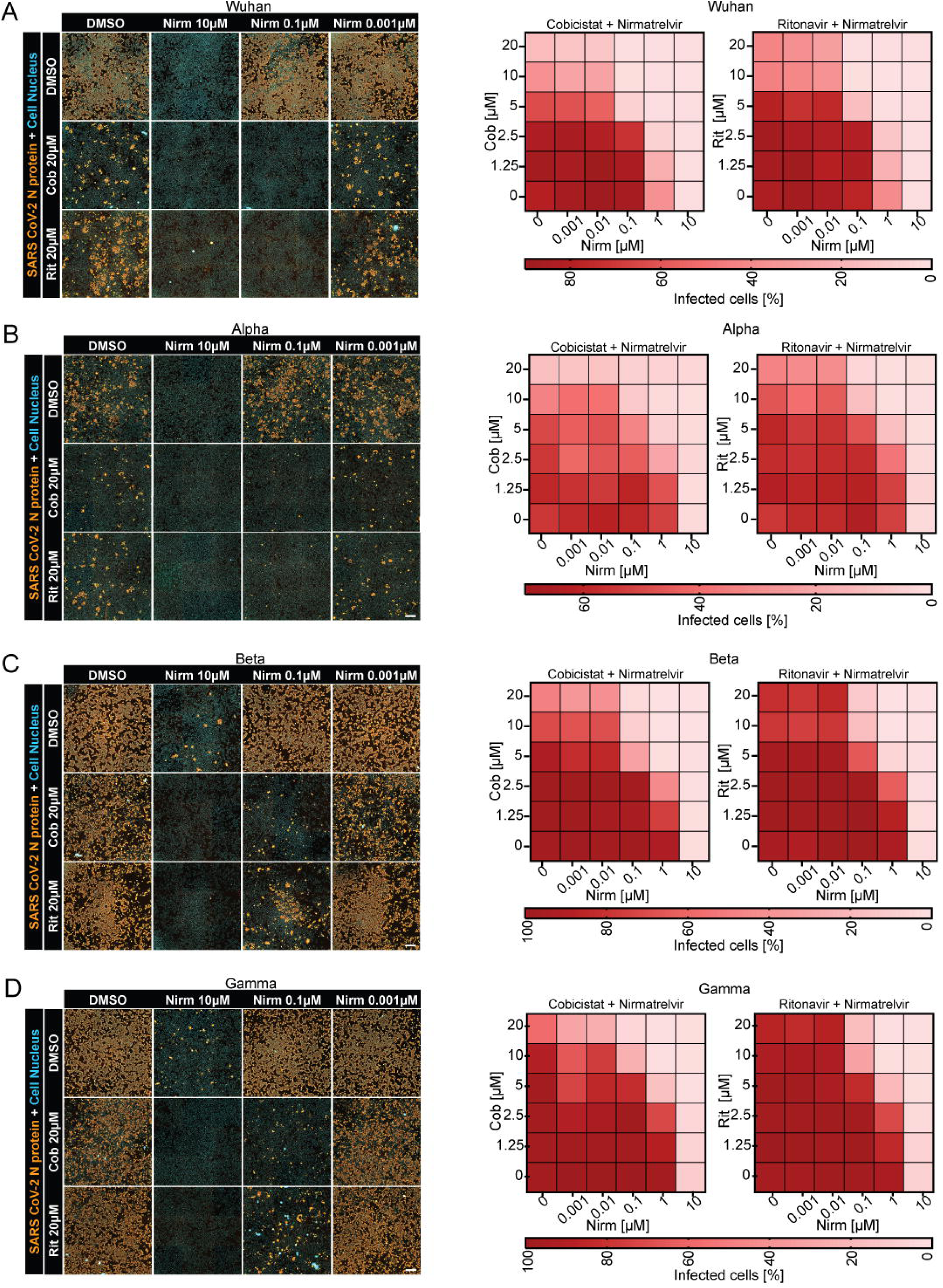
Combined antiviral activity of nirmatrelvir with cobicistat or ritonavir against the reference Wuhan SARS-CoV-2 strain and regionally spread VOCs. Panels A-D. VTN cells were pre-treated with cobicistat or ritonavir, alone or in combination with nirmatrelvir, at the indicated concentrations. Cells were then then infected with the Wuhan (A), Alpha (B), Beta (C) or Gamma (D) variants. After 24h the cells were fixed and stained for DAPI (blue) and the viral N protein (orange). Representative images are shown in the left panels. The percentage of cells positive for the N protein were then plotted as dose-dependent matrix heatmaps for each SARS-CoV-2 variant. The heatmaps depict mean values of two independent experiments, each performed in duplicate (right panels). Scale bar = 500 μm.

**Figure 5.**
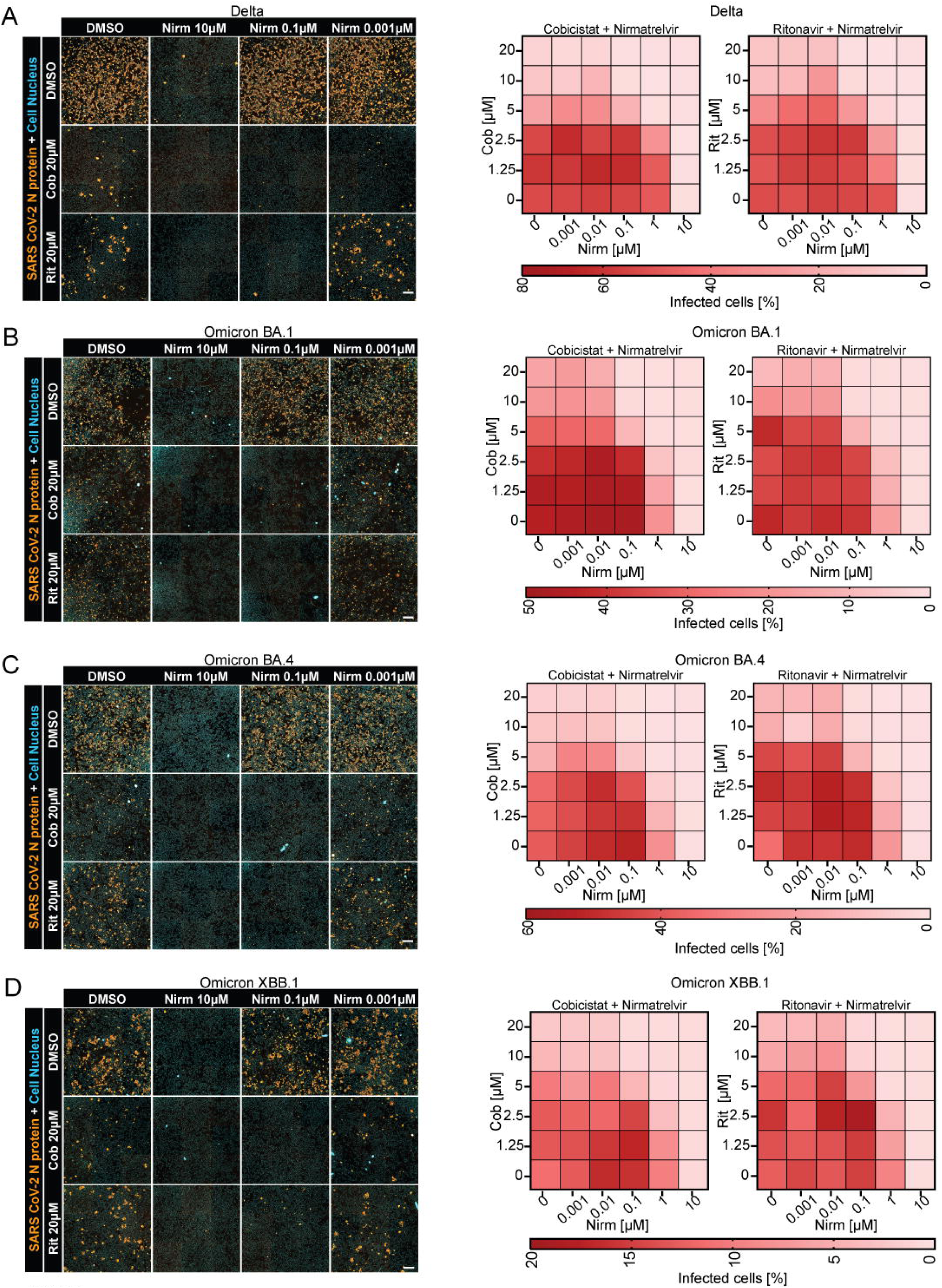
Combined antiviral activity of nirmatrelvir with cobicistat or ritonavir against SARS-CoV-2 VOCs and VOIs characterized by current or previous global spread. Panels A-D. VTN cells were pre-treated with cobicistat or ritonavir, alone or in combination with nirmatrelvir, at the indicated concentrations. Cells were then then infected with the Delta (A), Omicron BA.1 (B), Omicron BA.4 (C) or Omicron XBB.1 (D) SARS-CoV-2 variants. After 24h cells were fixed and stained for DAPI (blue) and the viral N protein (orange). Representative images are shown in the left panels. The percentage of cells positive for the N protein were then plotted as dose-dependent matrix heatmaps for each SARS-CoV-2 variant. The heatmaps depict mean values of two independent experiments, each performed in duplicate (right panels). Scale bar = 500 μm.

To further dissect the type and strength of interaction of each drug combination we calculated parameters of synergy and potency using the SynergyFinder plus tool (Zheng et al., 2022). Synergy scores were estimated using four different methods (ZIP, Loewe, HSA, and Bliss) and both the cobicistat/nirmatrelvir and the ritonavir/nirmatrelvir combinations displayed similar, pan-variant, synergism (Figure 6, Figure S4) which was more evident in the highest range of the tested concentrations (5-20µM) of cobicistat and ritonavir (Figure S4). The potency of each drug combination was then analyzed by calculating combination sensitivity scores (CSS), as previously described (Malyutina et al., 2019). Of note, cobicistat/nirmatrelvir showed a significantly higher combination sensitivity score than ritonavir/nirmatrelvir (Figure 6), suggesting enhanced suppression of SARS-CoV-2 replication across the range of concentrations tested, and in line with the increased potency of single treatment with cobicistat (*i.e.* relative inhibition of CYP3A inhibitors alone, RI) (Figure 6).

**Figure 6.**
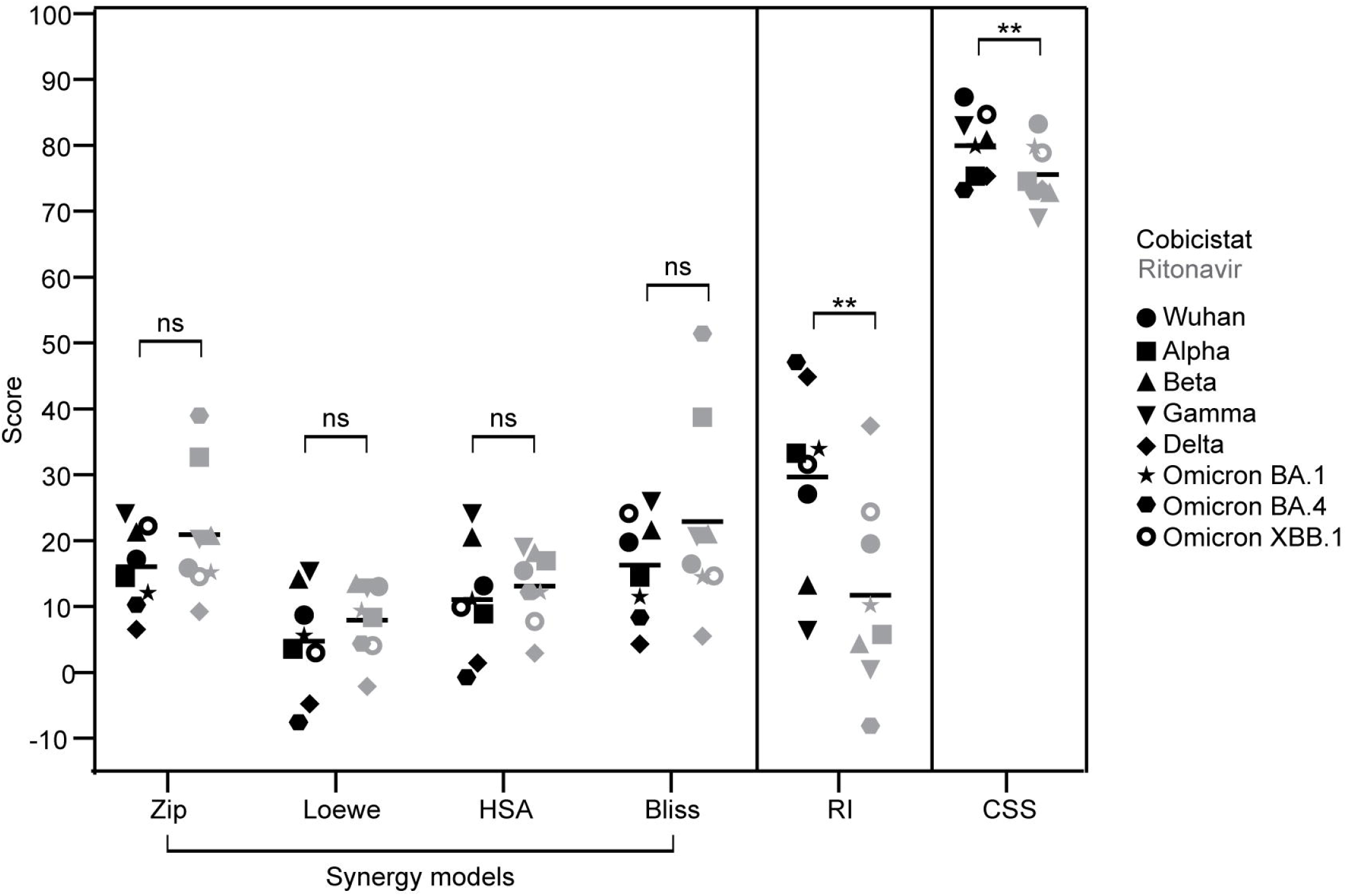
Comparison of the synergy and efficacy indices of the cobicistat/nirmatrelvir and the ritonavir/nirmatrelvir combinations. The data shown in Figures 4 and 5 were used to calculate the relative inhibition of viral replication induced by each drug or drug combination, as compared to vehicle (DMSO) controls. Inhibition levels were used to compute synergy (ZIP, Loewe, HSA, and Bliss models), relative inhibition (RI) by single treatment with cobicistat or ritonavir and combination sensitivity scores (CSS) (Malyutina et al., 2019) using the SynergyFinder plus software (Zheng et al., 2022). The scores obtained with cobicistat (with nirmatrelvir) or ritonavir (with nirmatrelvir) were then compared by paired t-test. ** p <0.01.

Taken together, these results indicate that, although cobicistat and ritonavir have a similar synergistic interaction with nirmatrelvir, a more profound suppression of viral replication is obtained with the cobicistat/nirmatrelvir combination, likely reflecting the higher intrinsic anti-SARS-CoV-2 potency of cobicistat.

### 3.3. MERS-CoV and HCoV-229E are susceptible to inhibition by cobicistat and ritonavir

We finally investigated whether the antiviral properties of cobicistat and ritonavir could be extended to other pathogenic human coronaviruses. To this aim we used both an endemic, but mildly pathogenic (HCoV-229E), and a highly pathogenic (MERS-CoV) coronavirus species. Importantly, these viruses use different entry receptors as compared to SARS-CoV-2 (aminopeptidase N for HCoV-229E and dipeptidyl peptidase 4 for MERS-CoV) (Yeager et al., 1992; Raj et al., 2013). Therefore, these experiments also allowed to test whether the antiviral activity of CYP3A inhibitors, which (at least for cobicistat) is associated with viral fusion impairment (Shytaj et al., 2022), is dependent on ACE-2-mediated entry. We used Huh-7 and VTN cells for HCoV-229E and MERS-CoV infection, respectively, in line with their previously described susceptibility to infection (Matsuyama et al., 2020; Tang et al., 2005). Infection with HCoV-229E was assessed by staining for dsRNA while for MERS-CoV we quantified infection by staining specifically for its N protein.

In line with the results obtained on the SARS-CoV-2 VOCs and VOIs, both cobicistat and ritonavir were effective in inhibiting HCoV-229E (Figure 7A-C) and MERS-CoV (Figure 7 D-F) replication at low micromolar levels. While at the highest concentrations tested (*i.e.* 20 µM) both drugs also exhibited some cytotoxicity in Huh-7 cells (Figure S5), the antiviral effects were evident at well tolerated concentrations (i.e. 2.5-10 µM). For both coronaviruses, the EC50 concentrations of cobicistat were lower (Figures 7B,E). Although the difference was modest for MERS-CoV (Figure 7E), it reached statistical significance for HCoV-229E (Figure 7B), thus corroborating the more potent anti-coronavirus activity of cobicistat as compared to ritonavir.

**Figure 7.**
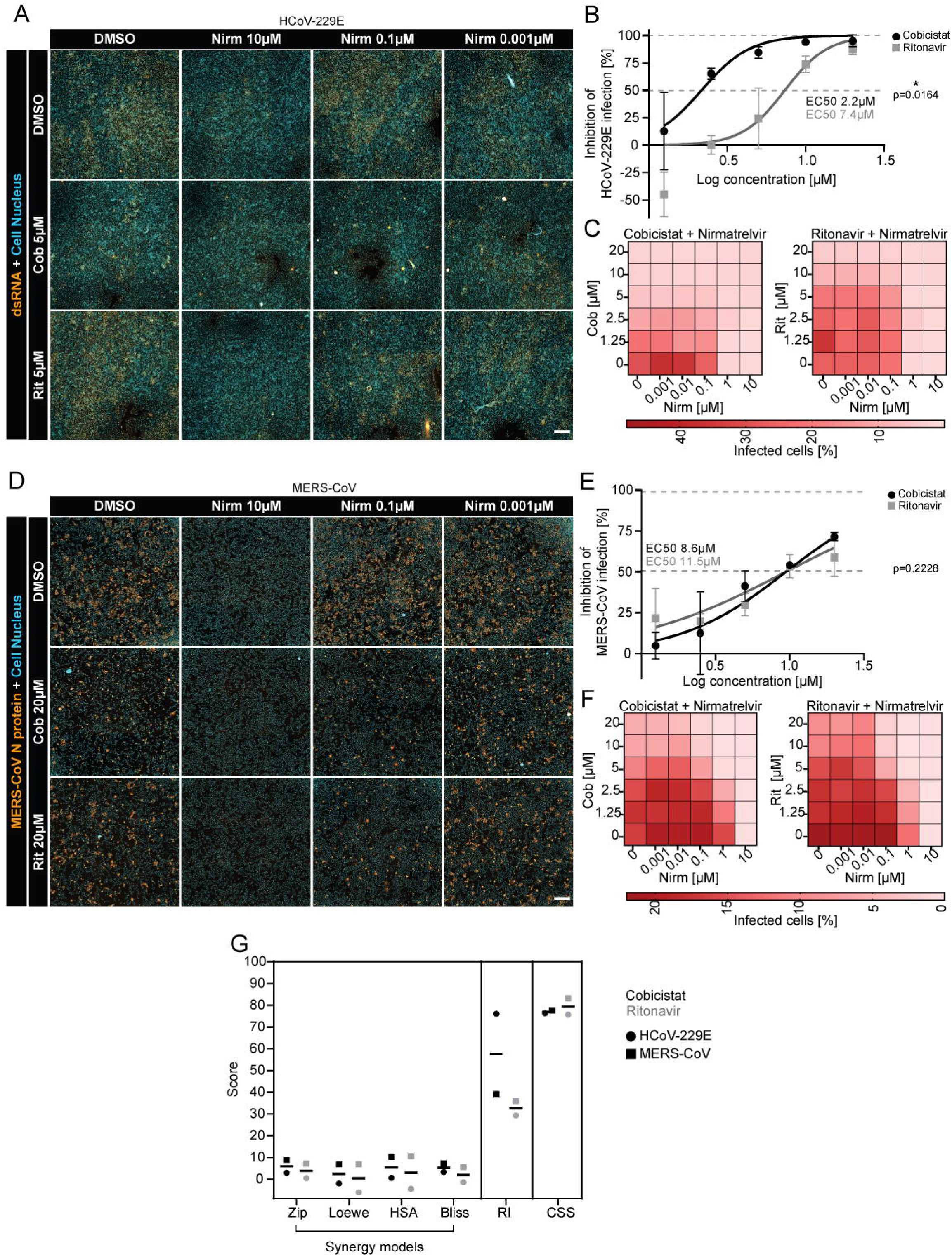
Susceptibility of HCoV-229E and MERS-CoV to inhibition by cobicistat or ritonavir, alone and in combination with nirmatrelvir. Panels A-C. Huh-7 cells were pre- treated with cobicistat or ritonavir, alone or in combination with nirmatrelvir, at the indicated concentrations. Cells were then infected with HCoV-229E and, after 24h, fixed and stained for DAPI (blue) and dsRNA (orange). Representative images are shown in panel A. The proportion of cells positive for dsRNA was calculated and used to derive the relative inhibition of viral replication as compared to vehicle (DMSO) controls. Half maximal effective concentrations (EC50) of both cobicistat and ritonavir were calculated by non-linear regression using a variable Hill slope (panel B). The percentage of infected cells upon treatment with drug combinations are depicted in dose-dependent matrix heatmaps (panel C). Panels D-F. VTN cells were pre-treated with cobicistat or ritonavir, alone or in combination with nirmatrelvir, at the indicated concentrations. Cells were then then infected with MERS-CoV and, after 24h, fixed and stained for DAPI (blue) and N protein (orange). Representative images are shown in panel D. The proportion of cells positive for the N protein was calculated and used to derive the relative inhibition of viral replication as compared to vehicle (DMSO) controls. Half maximal effective concentrations (EC50) of both cobicistat and ritonavir were calculated by non-linear regression using a variable Hill slope (panel E). The percentage of infected cells upon treatment with drug combinations are depicted in dose-dependent matrix heatmaps (panel F). Panel G. Viral inhibition levels were used to compute synergy (ZIP, Loewe, HSA, and Bliss models), relative inhibition by single treatment with cobicistat or ritonavir (RI), and combination sensitivity scores (CSS) (Malyutina *et al*., 2019) using the SynergyFinder plus software (Zheng *et al*., 2022). The divergence of the curves used to fit EC50 values for cobicistat and ritonavir was assessed by extra sum-of-squares F test (panels B and E). Each data point in panels B and E depicts mean ± SD of three independent experiments, each performed in duplicate. Panels C, F depict mean values of three independent experiments, each performed in duplicate. Scale bar = 500 μm. * p <0.05

We also tested the effects associated with the addition of nirmatrelvir, which is known to inhibit the replication of both viruses (Li et al., 2023; Owen et al., 2021). Although adding nirmatrelvir to either cobicistat or ritonavir was well-tolerated also in Huh-7 cells (Figure S5), surprisingly it did not lead to a synergistic inhibitory effect on HCoV-229E replication (Figure 7G). This was possibly a consequence of the higher potency of cobicistat on this virus, as suggested by the similarly high level of relative inhibition obtained with single treatment (RI score) as compared to combined treatment with cobicistat plus nirmatrelvir (CSS value) (Figure 7G). On the other hand, against MERS-CoV, combining nirmatrelvir with either cobicistat or ritonavir led to synergistic effects and comparable levels of overall inhibition of viral replication (Figure 7F,G). Altogether, these data reveal that cobicistat and ritonavir have a broad-spectrum anti-coronavirus activity and that the generally higher potency of cobicistat can be observed also against human coronaviruses other than SARS-CoV-2.

## 4. Discussion

The repurposing of clinically approved compounds for other indications is a powerful approach to accelerate drug discovery. This work shows that the antiviral activity of cobicistat, which we had previously demonstrated on two early circulating SARS-CoV-2 isolates (Shytaj et al., 2022), is conserved throughout multiple SARS-CoV-2 variants and coronavirus species. Our data also show that the parent drug of cobicistat, ritonavir, has similarly broad anti-coronavirus effects, but with consistently lower potency *in vitro*. Of note, the range of antiviral concentrations that we here observed on a broad panel of SARS-CoV-2 variants and pathogenic human coronaviruses are comparable with our previous results (Shytaj et al., 2022) obtained testing cobicistat on the early SARS-CoV-2 isolate Ger/BavPat1/2020 (EC50 4.1-7.7 µM in Vero E6 cells) and with independent studies testing ritonavir on the Wuhan SARS-CoV-2 variant (EC50 19.88µM) (Zhang et al., 2020) or using a recombinant MERS-CoV/luciferase construct (EC50 24.9µM) (Sheahan, Sims, Leist, et al., 2020).

The exact mechanism of action for the direct antiviral activity of cobicistat and ritonavir is still unclear. In particular, the lower potency of ritonavir is surprising given the close structural similarity between the two drugs (Xu et al., 2010) and the original design of cobicistat as a CYP3A inhibitor devoid of the anti-HIV protease activity of ritonavir (Xu et al., 2010). The similar potency of cobicistat and ritonavir against CYP3A, which is exerted at low nanomolar concentrations (Hossain et al., 2017; Xu et al., 2010), indicates that CYP3A inhibition is unlikely to play a significant role in the direct antiviral effects here described, which instead require low micromolar concentrations. Due to their flexibility in adapting to the catalytic site of several proteases both ritonavir and cobicistat were initially hypothesized to target the M^pro^ of SARS- CoV and SARS-CoV-2 (Savarino, 2005; Sharma et al., 2021). However, although the structure of cobicistat allows an initially favorable interaction with M^pro^, the calculated binding entropy suggests an unstable association and our previous experiments proved that cobicistat cannot inhibit this target (Shytaj et al., 2022).

By using cells overexpressing ACE-2 and the SARS-CoV-2 S protein, we previously showed that cobicistat can block SARS-CoV-2 fusion (Shytaj et al., 2022). The results herein obtained with HCoV-229E and MERS-CoV prove that the antiviral effects are independent of ACE-2, suggesting that cobicistat/ritonavir might inhibit viral entry through a conserved effect on the S protein. Interestingly, ritonavir is known to induce alterations of lipid-metabolism *in vitro* and *in vivo*, especially at higher dosages (Purnell et al., 2000; Riddle et al., 2001). Membrane cholesterol was shown to be essential for S protein mediated fusion and entry (Sanders et al., 2021; Wang et al., 2020) and lipids or lipid-modifying drugs have been proposed as broad-spectrum inhibitors of SARS-CoV-2 and other human coronaviruses (Toelzer et al., 2020; Wang et al., 2020). Our previous and current results, however, indicate that the antiviral effects of cobicistat are not enhanced by pre-treating cells before infection (Shytaj et al., 2022). This suggests that, if lipid alterations induced by cobicistat and ritonavir are associated with an effect on the S protein, this activity is exerted intracellularly and following the first cycle of viral entry. Future studies will be required to evaluate this hypothesis.

The synergistic effects that we observed when adding the M^pro^ inhibitor nirmatrelvir are in line with its known metabolism through CYP3A (Lamb, 2022), but peak synergy scores were detected at the highest concentrations of cobicistat and ritonavir, thus suggesting a contribution of the direct antiviral activity of CYP3A inhibitors. Due to the lack of consensus on how to appropriately quantify drug synergism, we compared and reported the results obtained with four (*i.e.* ZIP, Loewe, HSA, and Bliss) widely adopted synergy models (Greco et al., 1995; Zheng et al., 2022). We found similar synergy scores when nirmatrelvir was combined with cobicistat or ritonavir, in line with the comparable boosting activity previously described when these drugs are co-administered with HIV protease inhibitors or other CYP3A substrates (Tseng et al., 2017). While synergy provides an estimate of drug interactions, it does not necessarily capture therapeutic efficacy, which is arguably a more important parameter for real-world applications of drug combinations (Malyutina et al., 2019). Towards this purpose, we employed the previously described CSS parameter (Malyutina et al., 2019) and found that the cobicistat/nirmatrelvir combination induces a significantly more potent suppression of SARS-CoV-2 replication. On the other hand, CSS values for both combinations were similar when considering HCoV-229E and MERS-CoV. This difference might be in part explained by the fact that single treatment with cobicistat was *per se* able to completely suppress HCoV-229E replication at the concentration-range where interactions with nirmatrelvir are higher (5-20 µM).

The direct anti-coronavirus activity of cobicistat and ritonavir is only detectable at low micromolar concentrations. Based on our experiments, the ideal concentration range would be 5- 10 µM for cobicistat which, in combination with 0.1-1 µM nirmatrelvir, was both well tolerated and associated with complete (or almost complete) inhibition of all tested coronaviruses. Leveraging these antiviral properties would therefore require higher dosing regimens than those used in the Paxlovid formulation (Lamb, 2022), in typical antiretroviral combinations for people living with HIV (Tseng et al., 2017) and in clinical trials attempted in the early stages of the SARS-CoV-2 pandemic (Cao et al., 2020; Chen et al., 2020). In this regard, previous data obtained treating SARS-CoV-2 infected Syrian hamsters (Shytaj et al., 2022) as well as pharmacokinetic studies in uninfected mice (Pharmacology Review of Cobicistat, New Drug Application nr. 203-094) suggest that these concentrations could be achievable and relatively safe, at least for short-term administration against acute infection. It is however important to note that the profile of interactions of cobicistat and ritonavir with other drugs, including commonly prescribed medications (Tseng et al., 2017), might change at higher dosages due to their increasing affinity for CYPs other than CYP3A (Xu et al., 2010). Clinical dose escalation studies will be required to evaluate the safety and feasibility of this approach in humans.

The main limitation of our study is the lack of a mechanistic explanation of the different antiviral potencies of cobicistat and ritonavir, although our data indicate that both drugs target a conserved step in the life cycle of human coronaviruses. Moreover, even though previous literature reports comparable plasma concentrations upon equivalent dosing of cobicistat or ritonavir (Hsu et al., 1997; Mathias et al., 2010) a study comparing the antiviral effects of the two drugs *in vivo* will be required to confirm our *in vitro* finding of the higher potency of cobicistat.

Overall, our study shows that the CYP3A inhibitors ritonavir and, to a higher extent, cobicistat can be repurposed as broadly effective anti-coronavirus agents at concentrations potentially achievable *in vivo* by adjusting currently approved dosing regimens.

## Acknowledgments

ILS acknowledges Institutional funding from the Faculty of Life Sciences (University of Bristol, grant code U102054-101). ADD is a member of the G2P-UK National Virology consortium funded by the Medical Research Council/UKRI (Grant MR/W005611/1) that supplied SARS-CoV-2 variants. The authors acknowledge support from the University of Bristol’s Alumni and Friends, who funded the ImageXpress Pico Imaging System. The authors thank Dr. Irene Carlon-Andres and Dr. David Williamson (King’s College London, UK) for helpful suggestions on Fiji macro coding and Dr. Andrea Savarino (Italian Institute of Health, Italy) for critical reading of the manuscript.

## Supplementary Figures

**Figure S1. Effects of cobicistat or ritonavir on the viability of VTN cells.** VTN cells were treated with either cobicistat or ritonavir at the same concentrations used for antiviral assays (Figures 1,2) and their viability was assessed by MTT assay 24h post-treatment. The results are expressed as relative viability normalized to vehicle (DMSO) controls, with each data point showing the mean ± SD of three experiments, each performed in duplicate.

**Figure S2. Staining and automated image analysis of SARS-CoV-2 infected cells.** Panel A. Representative images of uninfected and SARS-CoV-2 infected VTN cells following immunofluorescent staining for the N viral protein (orange) and staining for nuclei (DAPI, in blue). Scale bar = 500 μm. Panel B. Automated segmentation and identification of nuclei negative (violet) or positive (white) for the SARS-CoV-2 N protein. Scale bar = 50 μm. All images were acquired and analyzed using the ImageXpress Pico Automated Cell Imaging System platform and software.

**Figure S3. Effect of the addition of nirmatrelvir to cobicistat or ritonavir on the viability of VTN cells.** VTN cells were treated with either cobicistat or ritonavir, alone or in combination with nirmatrelvir, at the same concentrations used for antiviral assays (Figures 3 and 4) and their viability was assessed by MTT assay 24h post-treatment. The heatmap shows the relative viability normalized to vehicle (DMSO) controls, with each data point depicting the mean of three independent experiments, each performed in duplicate.

**Fig. S4. Representative synergy 3D plots of the interaction between cobicistat or ritonavir with nirmatrelvir.** Synergy plots were generated with the SynergyFinder plus software (Zheng et al., 2022) using the data in Figures 4 and 5 to calculate the inhibitory effects of each drug combination (upper panel: cobicistat/nirmatrelvir; lower panel: ritonavir/nirmatrelvir). The Figure shows representative plots for the Wuhan (A), Gamma (B), and Omicron XBB.1 (C) SARS-CoV-2 variants. For each variant, four different synergy models (*i.e.* ZIP, Loewe, HSA, and Bliss) were used to calculate synergy scores and generate the plots.

**Fig. S5. Effect of cobicistat or ritonavir, alone or in combination with nirmatrelvir, on the viability of Huh-7 cells.** Huh-7 cells were treated with either cobicistat or ritonavir, alone or in combination with nirmatrelvir, at the same concentrations used for antiviral assays (Figure 7) and their viability was assessed by MTT assay 24h post-treatment. The heatmap shows the relative viability normalized to vehicle (DMSO) controls, with each data point depicting the mean of three independent experiments, each performed in duplicate.

