## Supplementary figures and images for "Broad-spectrum antiviral activity of clinically approved CYP3A inhibitors against pathogenic human coronaviruses in vitro"

### Supplementary Figure 1

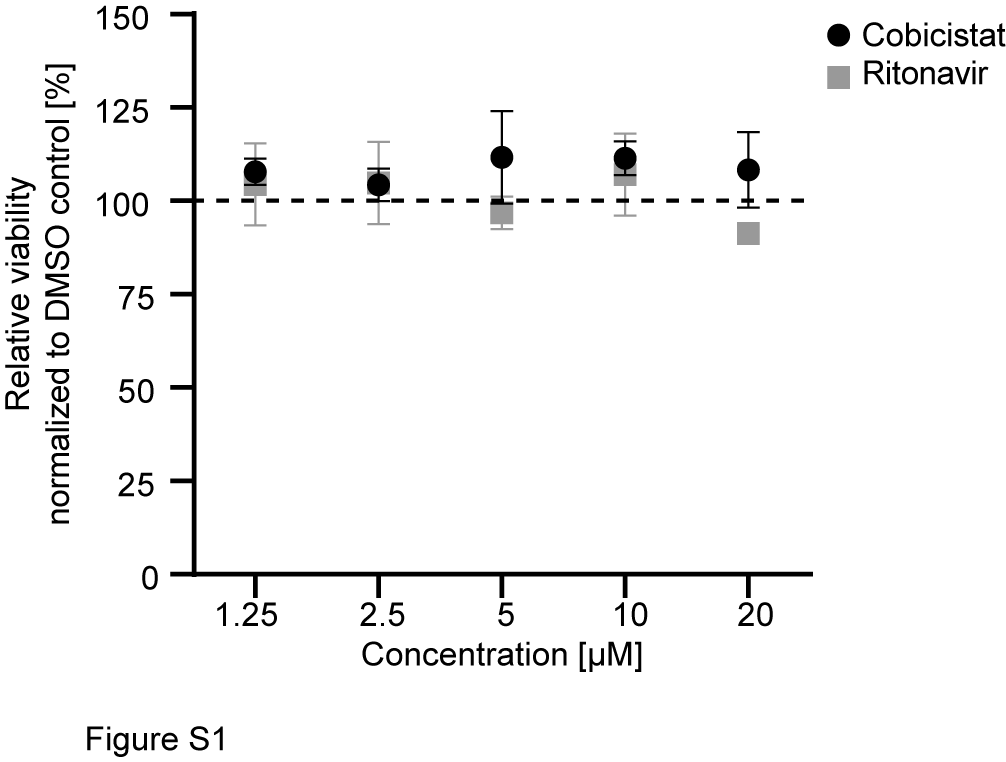

### Supplementary Figure 2

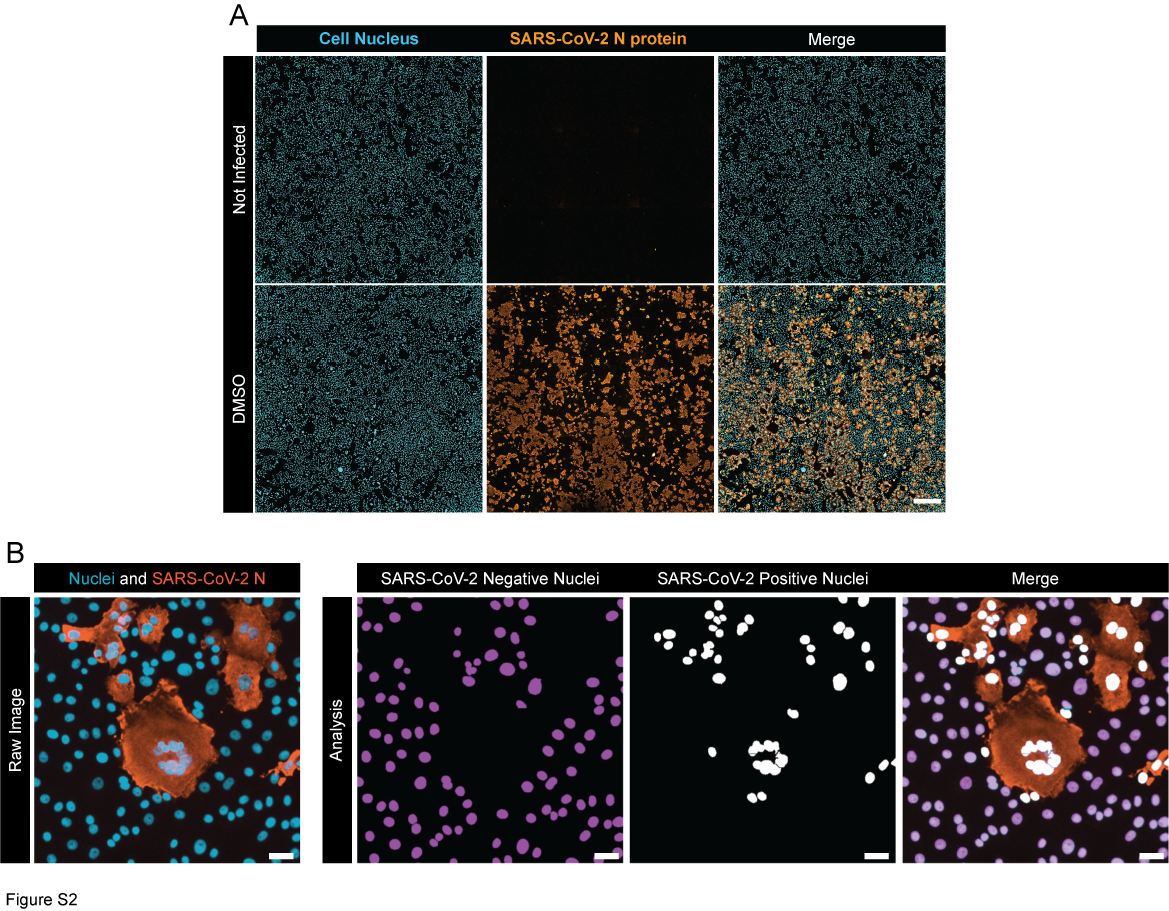

### Supplementary Figure 3

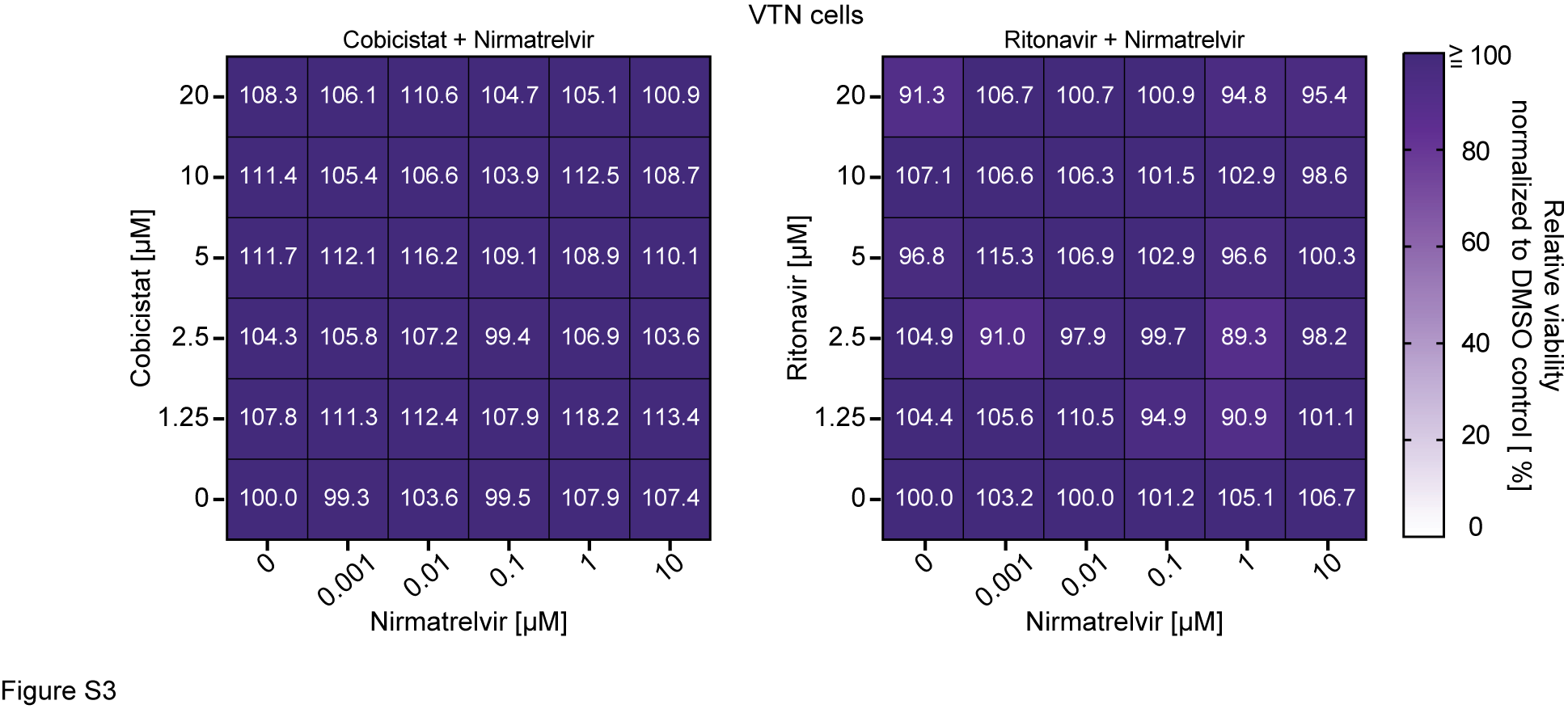

### Supplementary Figure 4

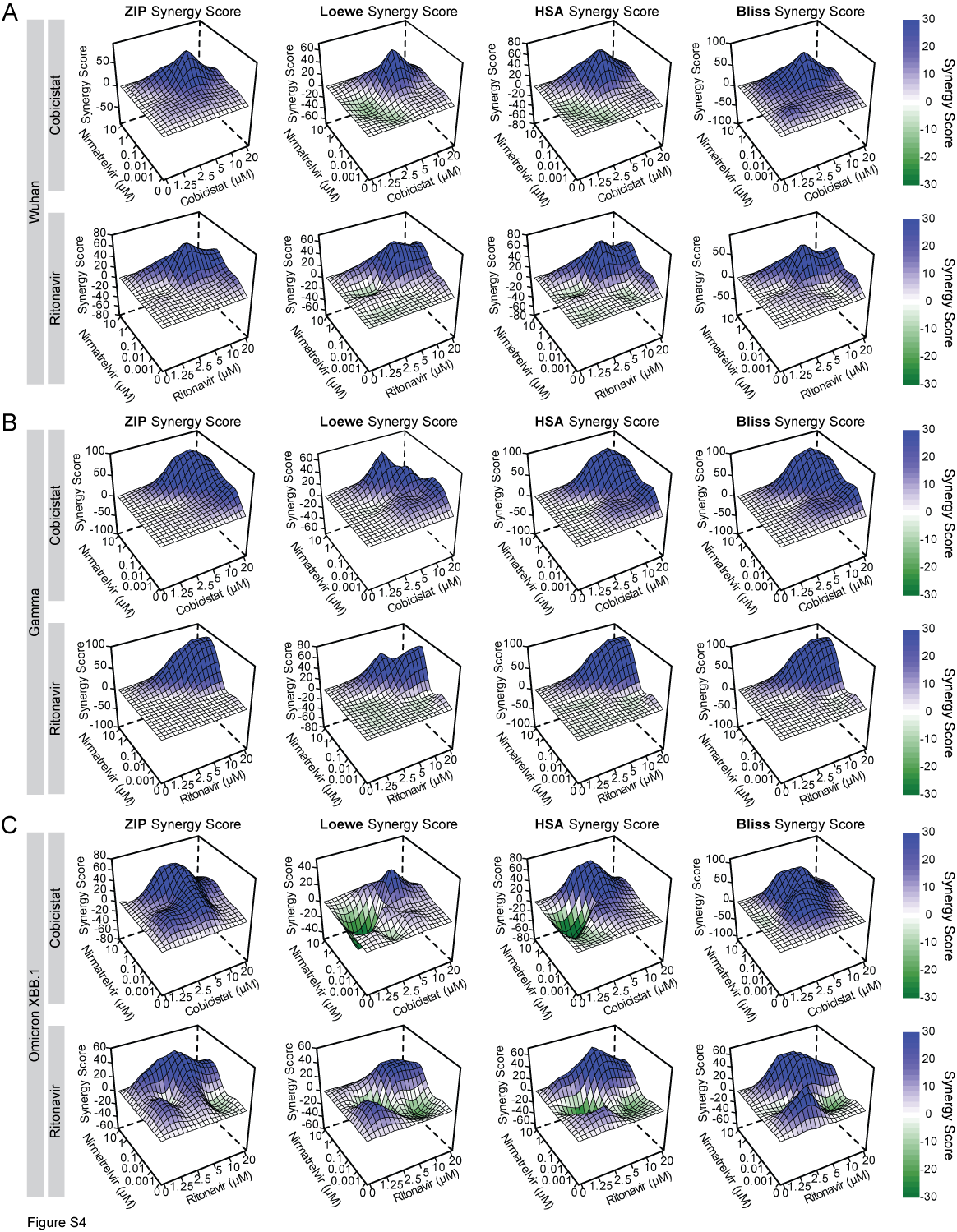

### Supplementary Figure 5

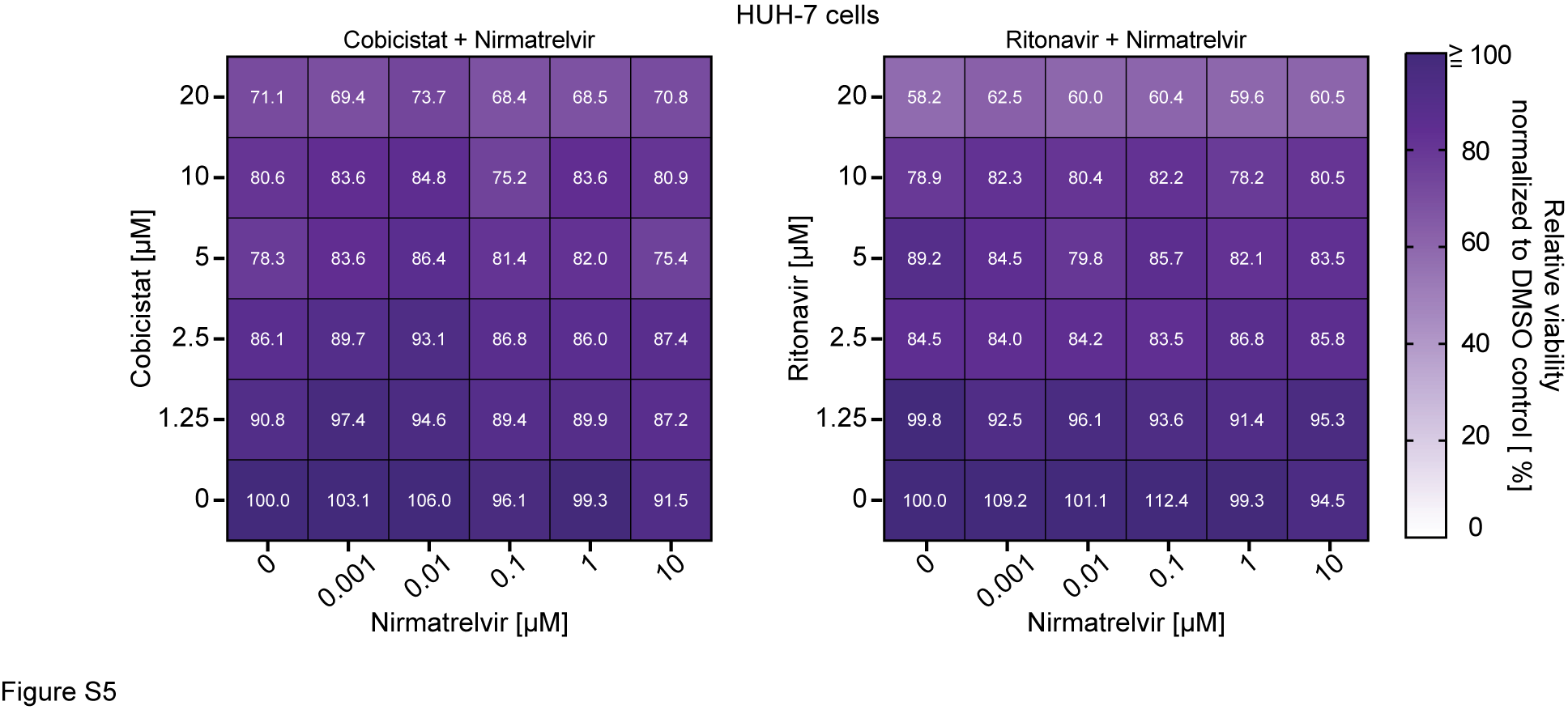
